# Repression of divergent noncoding transcription by a sequence-specific transcription factor

**DOI:** 10.1101/314310

**Authors:** Andrew CK Wu, Harshil Patel, Minghao Chia, Fabien Moretto, David Frith, Ambrosius P. Snijders, Folkert van Werven

## Abstract

Many active eukaryotic gene promoters exhibit divergent noncoding transcription, but the mechanisms restricting expression of these transcripts are not well understood. Here we demonstrate how a sequence-specific transcription factor represses divergent noncoding transcription at highly expressed genes in yeast. We find that depletion of the transcription factor Rap1 induces noncoding transcription in a large fraction of Rap1 regulated gene promoters. Specifically, Rap1 prevents transcription initiation at cryptic promoters near its binding sites, which is uncoupled from transcription regulation in the protein-coding direction. We further provide evidence that Rap1 acts independently of chromatin-based mechanisms to repress cryptic or divergent transcription. Finally, we show that divergent transcription in the absence of Rap1 is elicited by the RSC chromatin remodeller. We propose that a sequence-specific transcription factor limits access of basal transcription machinery to regulatory elements and adjacent sequences that act as divergent cryptic promoters, thereby providing directionality towards productive transcription.

## Introduction

Precise control of gene expression is of critical importance for all cellular functions across species. How and when genomes produce coding messenger RNAs and prevent the expression of unwanted noncoding RNAs has been a long-standing question of interest. In this context an apparent paradox exists: genomic locations of coding gene transcription also produce aberrant noncoding transcripts. A major source are the transcriptionally active coding gene promoters, which often express noncoding transcripts in the antisense direction (Neil et al., 2009; Seila et al., 2008; Sigova et al., 2013; Xu et al., 2009). This process is known as divergent or bidirectional transcription. The functions of the noncoding RNAs produced and the mechanisms that limit expression of divergent noncoding transcripts are not well understood.

Divergent noncoding transcription is present across eukaryotic species (Seila et al., 2009). A large fraction of all noncoding transcripts emanate from divergent or bidirectional gene promoters (Neil et al., 2009; Seila et al., 2008; Xu et al., 2009). Typically, divergent noncoding transcripts are initiated within or nearby coding gene promoters but they do not share the same core promoter sequence as transcripts in the coding direction (Andersson et al., 2015; Duttke et al., 2015; Rhee and Pugh, 2012; Scruggs et al., 2015). The transcription of divergent noncoding RNAs is generally lower than that of coding genes (Churchman and Weissman, 2011). Divergent noncoding transcripts are generally unstable and rapidly degraded (Neil et al., 2009; van Dijk et al., 2011). The Nrd1-Nab3-Sen1 (NNS) and premature polyadenylation signal (PAS) pathways in yeast and mammalian cells, respectively, terminate and degrade divergent transcripts using specific sequence elements (Almada et al., 2013; Arigo et al., 2006; Ntini et al., 2013; Schulz et al., 2013; Thiebaut et al., 2006). In addition, exosome and nonsense mediated decay pathways regulate RNA turnover and limit expression of cryptic transcripts originating from divergent promoters (Neil et al., 2009; van Dijk et al., 2011; Wery et al., 2016). Finally, a chromatin based-mechanism that limits divergent noncoding transcription at promoters has been identified (Marquardt et al., 2014). Specifically, CAF-1 mediated chromatin assembly represses the accumulation of divergent noncoding transcripts, which in turn is opposed by chromatin regulators that promote rapid turnover of nucleosomes.

In budding yeast, 138 genes encode for the protein subunits of the ribosome. These so-called ribosomal protein (RP) genes are highly transcribed and account for approximately half of all RNA polymerase II transcription in rapidly dividing cells (Li et al., 1999; Warner, 1999). Transcription of nearly all RP genes is controlled by the pioneer transcription factor Rap1, which binds to upstream sequence elements in RP gene promoters (Lieb et al., 2001). When RP gene promoters are active Rap1 recruits coactivators such as Fhl1, Ifh1, and Sfp1, as well as basal transcription factors like TFIID and TFIIA (Garbett et al., 2007; Hall et al., 2006; Marion et al., 2004; Papai et al., 2010; Reja et al., 2015; Rudra et al., 2005; Wade et al., 2004; Zhao et al., 2006). Thus, Rap1 orchestrates RP gene expression. Given that RP genes are among the most actively transcribed genes in yeast, they are an ideal model for studying how aberrant or cryptic divergent transcription is repressed at highly expressed gene promoters.

Here we describe how divergent noncoding transcription is repressed at the highly active ribosomal protein gene promoters. We find that depletion of Rap1, but not other transcription factors important for ribosomal protein expression, causes transcription in the divergent direction. Rap1 represses noncoding transcription typically within 50 base pairs of the Rap1 motif, which is uncoupled from transcription regulation in the protein-coding direction. We further show that Rap1 mediated repression of divergent transcription is distinct from chromatin based mechanisms. Thus, a sequence-specific transcription factor controls promoter directionality by repressing transcription in the divergent direction. Our work adds a new layer of regulation to the various mechanisms that limit expression of aberrant transcripts and defines how promoter directionality is controlled.

## Results

### Depletion of Rap1 causes divergent transcription at *RPL43B* and *RPL40B*

In budding yeast, a large fraction of bidirectional promoters express noncoding transcripts, also known as cryptic unstable transcripts (CUTs) or stable unannotated transcripts (SUTs), in the divergent direction (Neil et al., 2009; Xu et al., 2009). Transcription of divergent CUTs and SUTs typically correlates with nucleosome depleted regions (NDRs) and promoter activity in the coding gene direction (Xu et al., 2009). Considering that RP genes are among the most highly expressed genes in yeast, surprisingly few RP gene promoters (16 out of 138 promoters) display an annotated divergent noncoding transcript (CUT or SUT) (Xu et al., 2009). We hypothesized that RP promoters must have a robust mechanism for limiting divergent noncoding transcription. To investigate this, we assessed how depleting or deleting transcription factors important for RP gene regulation affects divergent transcription. We selected the *RPL43B* and *RPL40B* genes to study, since both promoters are directly adjacent to an annotated divergent noncoding transcript: *IRT2* and *SUT242*, respectively (Figure 1A). Four of the transcription factors (Fhl1, Ifh1, Sfp1, and Rap1) have an essential role in cellular fitness (Blumberg and Silver, 1991; Cherel and Thuriaux, 1995; Hermann-Le Denmat et al., 1994; Shore and Nasmyth, 1987). Hence, we generated depletion alleles by tagging the carboxy termini with the auxin inducible degron (AID) and expressed the TIR1 E3 ubiquitin ligase that targets AID for degradation in the presence of Indole-3-acetic acid (IAA) (Nishimura et al., 2009) (Figure 1B). We measured the expression of divergent transcripts by northern blot using probes directed against *IRT2* and *SUT242*. No effects on *IRT2* and *SUT242* expression were observed when we depleted Fhl1, Ifh1, or Sfp1, or in *hmo1Δ* or *crf1Δ* cells (Figure 1C). Strikingly, Rap1 depleted cells (*RAP1-AID* + IAA) showed strong induction of a transcript resembling *IRT2* (Figure 1C). In addition, the *RPL40B* promoter displayed expression of multiple divergent transcripts upon Rap1 depletion. The transcript with the strongest signal approximated the size of the adjacent *MLP1* gene, which we define as isoform of *MLP1* (*iMLP1*)*. IRT2* and *iMLP1* expression increased simultaneously as Rap1 protein levels decreased when we followed the kinetics of divergent transcription (Figure 1D and Figure S1A). These data show that Rap1, but not the other transcription factors important for RP expression, represses divergent transcription at the *RPL43B* and *RPL40B* promoters.

**Figure 1.**
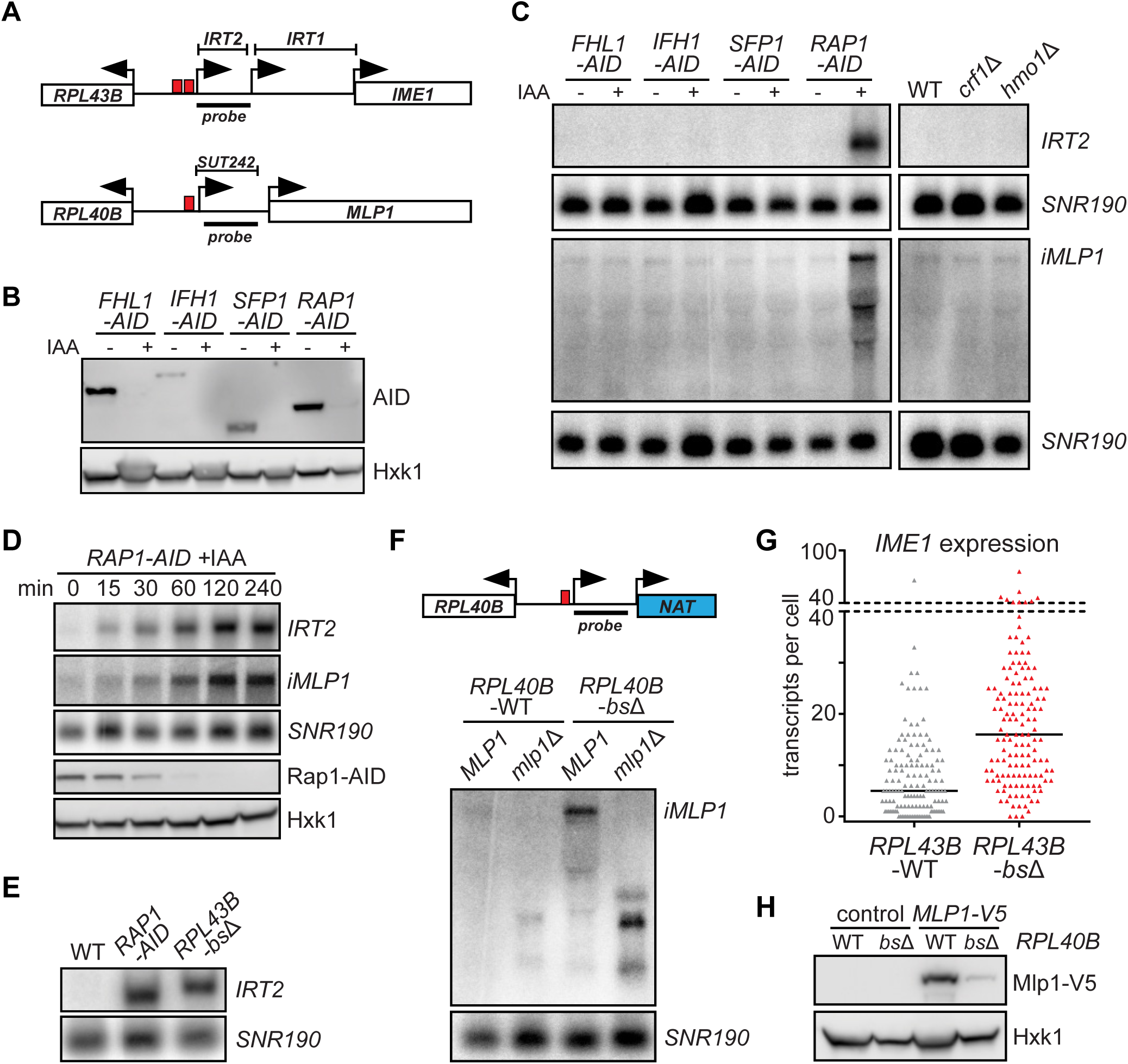
Rap1 prevents expression of noncoding RNAs. **(A)** Schematic overview of two divergent ribosomal protein gene promoters. **(B)** Auxin induced depletion (AID) of transcription factors important for ribosomal protein gene expression. Transcription factors were tagged at the carboxy-terminus with the auxin inducible degron to generate: *FHL1-AID* (FW4200), *IFH1-AID* (FW4202), *SFP1-AID* (FW4204), and *RAP1-AID* (FW3877). Cells were grown to exponential phase, either not treated (0 hours) or treated with 3-indole-acetic acid (IAA, 500 μM) for 2 hours. Protein extracts were separated by SDS-PAGE, blotted, and AID-tagged proteins were detected with an anti-V5 antibody. Hxk1 was used as a loading control. **(C)** *IRT2* and *iMLP1* expression in cells described in A. For the northern blot analyses wild-type control (FW629), *crf1Δ* (FW4136), and *hmo1Δ* (FW4132) cells were also included. RNA was extracted, separated by gel electrophoresis, blotted, and membranes were hybridized with a 32P-labelled probes targeting *IRT2* or *SUT242*, and *SNR190* as a loading control. **(D)** Similar as B and C, except that *RAP1-AID* (FW3877) cells were used for analysis in a time course. **(E)** *IRT2* expression in *RPL43B-bsΔ* cells (FW3443), which harbour a deletion of the Rap1 binding sites in the *RPL43B* promoter. For the analyses wild-type (FW629), and Rap1 depleted (FW3877) cells were also included as described in C. **(F)** Similar as E, except that *iMLP1* expression in *RPL40B-bsΔ* (FW4141) cells, which harbour a deletion of the Rap1 binding site in the *RPL40B* promoter, *mlp1Δ* cells (FW6030) and *mlp1Δ RPL40B-bsΔ* (FW6029) cells. **(G)** Distribution of *IME1* expression in single diploid S288C cells, with intact Rap1 binding sites in the *RPL43B* promoter (FW631) or Rap1 binding sites deleted (*RPL43B-bsΔ*, FW6139). Cells were grown in rich medium (YPD) until saturation and subsequently shifted to sporulation medium (SPO). Samples were fixed, and hybridized with fluorescent-labelled probes directed against *IME1* (AF594) and *ACT1* (Cy5) mRNAs prior to imaging. Each triangle represents the transcript count for one cell and black lines indicate median number of transcripts per cell. For the analysis n = 139 cells were used. *p < 0.0001 (unpaired student’s t-test). **(H)** Mlp1-V5 expression in the presence and absence of Rap1 binding sites in the *RPL40B* promoter. Wild-type (FW629), *RPL40B-bsΔ* (FW4141), or *MLP1* tagged with V5 epitope tag (*MLP1-V5*, FW4122) and *MLP1-V5 RPL40B-bsΔ* (FW4120) strains were used for the analysis.

Rap1 is a pioneer transcription factor that binds to well-defined DNA sequence elements of RP and metabolic gene promoters (Azad and Tomar, 2016; Lieb et al., 2001; Zaret and Carroll, 2011). To examine whether the Rap1 binding site is important for repressing divergent noncoding transcription, we deleted Rap1 sequence motifs in the *RPL43B* and *RPL40B* promoters *(RPL43B-bsΔ* and *RPL40B-bsΔ*)*. IRT2* expression levels increased in *RPL43B-bsΔ* cells, to a level comparable to Rap1 depleted cells (*RAP1-AID* +IAA) (Figure 1E). It is worth noting that initiation of *IRT2* transcription occurred downstream of the Rap1 sites in *RPL43B-bsΔ* because the loxP sequence (that was used to generate *RPL43B-bsΔ*) increased the *IRT2* transcript length (Figure 1E and Figure S1B). The expression of *iMLP1* also increased in cells lacking the Rap1 binding site in the *RPL40B* promoter (*RPL40B-bsΔ*) (Figure 1F, compare lane 1 to 3, and Figure S1B). Thus, Rap1 binding is required to repress divergent noncoding transcription from the *RPL43B* and *RPL40B* promoters.

It is well established that transcription within intergenic regions can affect local coding gene expression through transcriptional interference (Ard et al., 2017). This prompted us to examine the effect of divergent transcription on the expression of neighbouring genes at the *RPL43B* and *RPL40B* promoters. Previous work has shown that *IRT2* is part of a regulatory circuit that facilitates expression of *IME1*, (Moretto et al., 2018). In wild-type cells a Ume6 binding site, which is localized directly adjacent to the Rap1 motifs towards *IME1*, controls *IRT2* expression (Moretto et al., 2018). We hypothesized that Rap1 prevents mis-expression of *IRT2* from affecting *IME1* levels. When we measured *IME1* expression levels in single diploid cells during entry into meiosis, the median *IME1* expression increased from 5 transcripts per cell for the control *(RPL43B-WT)* to 16 transcripts per cell in the *RPL43B-bsΔ* mutant (Figure 1G and Figure S1C). We also investigated the effect of divergent transcription from the *RPL40B* promoter on *MLP1* expression. First, we deleted *MLP1* in the *RPL40B-bsΔ* background *(RPL40B-bsΔ mlp1*Δ), and found that the *iMLP1* transcript disappeared and a shorter transcript appeared, demonstrating that *iMLP1* is a long transcript isoform (Figure 1F). The 5’extended sequence of *iMLP1* harbours 15 upstream AUG sequences, 10 of which are out-of-frame, suggesting that the transcript is unlikely to allow translation of full-length Mlp1 protein similarly to other long undecodable transcript isoforms described previously (Chen et al., 2017; Cheng et al., 2018; Chia et al., 2017). Mlp1-V5 protein levels were markedly reduced in *RPL40B-bsΔ* compared to *RPL40B-WT* cells, suggesting that *iMLP1* transcription affects expression of the coding *MLP1* mRNA (Figure 1H and Figure S1D). We conclude that mis-regulation of Rap1-repressed divergent transcripts affects neighbouring gene expression.

### Rap1 represses noncoding transcription near its binding site

Having established that Rap1 is essential for preventing divergent transcription at the *RPL43B* and *RPL40B* gene promoters, we next investigated how depleting Rap1 affects noncoding transcription at a genome-wide scale by RNA sequencing (RNA-seq). To ensure detection of lowly expressed RNA Polymerase II transcripts, we performed RNA-seq on both polyadenylated RNA (polyA) and total RNA after ribosomal RNA depletion to sufficient depth (~45 million reads). As expected, the expression of Rap1 regulated coding genes decreased upon Rap1 depletion (compare DMSO to IAA) (Figure S2A and S2B) (Knight et al., 2014; Lieb et al., 2001). In addition, *IRT2* expression increased in IAA treated *RAP1-AID* cells, whereas the control (DMSO) did not show *IRT2* expression (Figure 2A). We also observed noncoding transcription from other RP gene promoters after Rap1 depletion. For example, the *RPL8A* promoter expressed a divergent transcript that spans the neighbouring *GUT1* gene, but antisense to the coding sequence (Figure 2A). Consequently, sense *GUT1* expression was reduced. Thus, RNA-seq is able to identify novel Rap1-repressed divergent transcripts.

**Figure 2.**
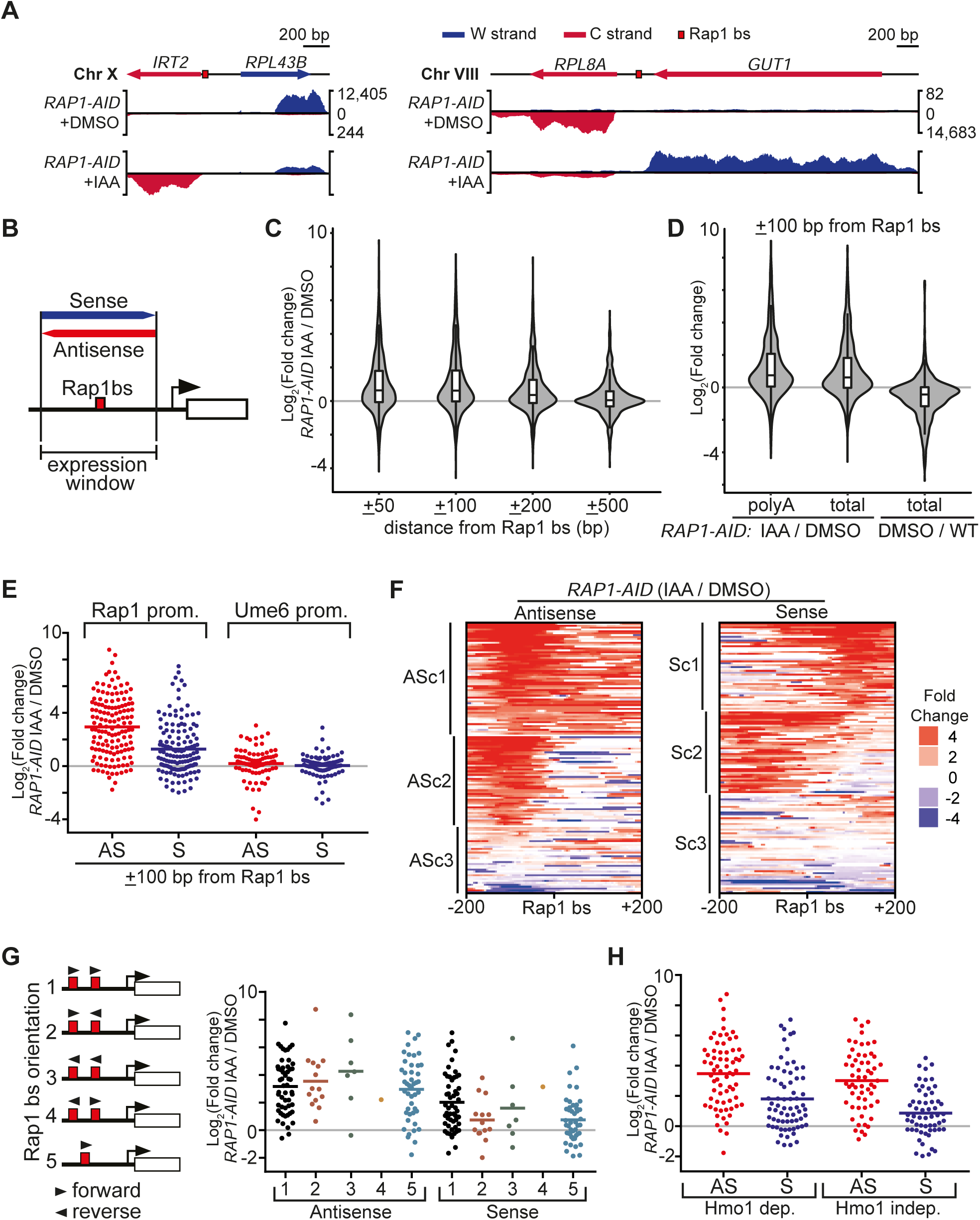
Rap1 represses divergent noncoding transcription. **(A)** Genome browser tracks showing examples of divergent noncoding RNAs repressed by Rap1. *RAP1-AID* cells (FW3877) were grown to exponential phase and were treated with DMSO or IAA for 2 hours. Samples were taken and processed for total RNA-seq. The normalized reads are shown on the y-axis for the Watson (W, blue) and Crick (C, red) strands. **(B)** Scheme for determining the RNA-seq signals around Rap1 motifs. The RNA-seq signals for specified distances up- and downstream of Rap1 binding sites were computed by taking the reads that overlapped with the selected genomic region (window size). **(C)** Violin and box-and-whisker plots of total RNA-seq data described in A, displaying the expression changes for different windows sizes (x-axis) of *RAP1-AID* +IAA versus *RAP1-AID* +DMSO (y-axis). In total, n = 564 Rap1 sites were used for the analyses. Signal for the W and C strands was computed separately resulting in n = 1128 data points. **(D)** Similar as C, except that polyadenylated (polyA) and total RNA-seq data were compared for genomic regions of 100 bp up-and downstream of Rap1 sites (±100 bp window). As a control the expression changes in *RAP1-AID* +DMSO over wild-type (WT, FW629) are displayed (total RNA-seq). **(E)** Similar data as D, except that scatter plots were used to display the expression changes for the antisense and sense strands relative to the coding gene. For the analysis we used (n = 141) Rap1-regulated promoters and (n = 87) Ume6 regulated promoters. We approximated the Ume6 binding site as −250 bp upstream of the coding gene. Horizontal red or blue lines indicate mean values. **(F)** Heat maps showing the changes in RNA expression on the antisense and sense strands for data described in E. Promoters were clustered based on antisense (ASc1-3) or sense (Sc1-3) strand signals using k-means clustering (k = 3). **(G)** Scheme displaying the different classes of ribosomal protein gene promoters based on Rap1 binding site motif orientation (left). The Rap1 motifs (red boxes) and corresponding orientation (forward and reverse arrowheads) are shown. Scatter plots of data described in E, except that the data was separated by orientation of the Rap1 motifs (right). Number of promoters in each class: 1 (n = 53), 2 (n = 14), 3 (n = 7), 4 (n = 1), and 5 (n = 45). **(H)** Similar data as E, except that the data was divided in Hmo1 dependent (n = 69) and Hmo1 independent (n = 58) promoters.

Our data of example loci indicate that Rap1 mediates repression of noncoding transcription from cryptic promoters close to its binding site. To systematically determine how Rap1 depletion affects noncoding transcription, we binned RNA expression signals from the RNA-seq data (total RNA) in windows of 50, 100, 200, and 500 base pairs (bp) up- and downstream of 564 annotated Rap1 sites (Figure 2B) (Lieb et al., 2001; Rhee and Pugh, 2011). Strikingly, our analyses revealed that for the smaller windows (50 and 100 bp) approximately 40% of Rap1 binding sites displayed increased RNA expression (more than 2-fold) upon Rap1 depletion (Figure 2C). For the larger windows (200 and 500 bp) the number of Rap1 sites showing increased RNA expression (more than 2-fold) decreased to 30% and 16%, respectively, suggesting that the effects are spatially limited to regions harbouring Rap1 elements. Rap1-repressed noncoding transcripts are polyadenylated, because our analyses with different window sizes showed little difference between RNA-seq data from polyadenylated (polyA) RNA and total RNA (Figure 2D and Figure S2C). Taken together, these data show that Rap1 represses transcription near Rap1 binding sites across the genome.

Next, we analyzed the RNA-seq data to decipher important features of cryptic transcript repression by Rap1. First, we determined whether there is a bias for the orientation of Rap1 repressed transcripts. We selected 141 Rap1 binding sites from well-annotated gene promoters regulated by Rap1 (mostly RP genes) (Knight et al., 2014; Lieb et al., 2001). We found that expression near the Rap1 binding sites was upregulated in both the sense and antisense direction after Rap1 depletion, however the largest increase in expression was detected in the antisense direction (7.7-fold mean increase for the antisense strand, and 2.4-fold for the sense strand) (Figure 2E). A control set of promoters regulated by the meiotic transcriptional repressor, Ume6, was not affected by Rap1 depletion (McKnight et al., 2016). Second, we clustered the data centered on the Rap1 binding site, and identified clusters of promoters showing different changes in expression (Figure 2F, Figure S2D-E). While antisense clusters 1 and 2 (ASc1 and ASc2) both displayed increased expression upstream of the Rap1 binding site, ASc1 also showed a mild increase of antisense RNA expression downstream of the Rap1 site (Figure 2F). In ASc3 few changes in expression were observed. When we clustered for the sense direction signals, we observed that transcripts were up-regulated both up- and downstream of the Rap1 site (Sc1 and Sc2). Finally, we examined whether the transcripts induced upon Rap1 depletion were enriched for specific classes of RP gene promoters. Rap1 regulated RP gene promoters can be classified according to the orientation of Rap1 binding site motifs, and dependence on the high mobility group protein Hmo1 (Hall et al., 2006; Knight et al., 2014; Reja et al., 2015). We found that the orientation or number of Rap1 motifs had little effect on the level of antisense or sense expression after Rap1 depletion (Figure 2G). RP gene promoters regulated by Hmo1 displayed a comparable increase in expression around the Rap1 binding sites, versus promoters that do not depend on Hmo1 (Figure 2H). In conclusion, Rap1 represses transcription nearby its binding sites in the antisense direction, and to lesser extent the sense direction, independent of RP gene promoter architecture.

### A proximal Rap1 motif is required and sufficient to repress divergent transcription

Our results demonstrate that upon Rap1 depletion, noncoding transcription occurs nearby its binding site. If close proximity of the Rap1 binding site to the cryptic promoter sequence is important for transcriptional repression, then increasing the distance between the Rap1 motif and cryptic promoter should impair repression of noncoding transcription. To test this, we integrated a spacer sequence between the Rap1 motifs and the *RPL43B* core promoter, and measured the effect on *IRT2* expression. We found that Rap1 was not able to repress *IRT2* in the presence of a spacer sequence (Figure 3A and 3B). When we integrated a spacer sequence of 400 bp to replace Rap1 binding sites (bsΔS), *IRT2* expression was de-repressed and the size of *IRT2* increased indicating that the core promoter of *IRT2* is downstream of the spacer sequence (relative to *RPL43B)*. Strikingly, we observed a similar pattern, when we integrated the spacer directly downstream of the Rap1 binding site (S), relative to *RPL43B*. The spacer sequence did not affect the ability of Rap1 to associate with its motifs at *RPL43B* promoter (Figure 3C). In conclusion, the Rap1 binding site must be nearby the cryptic promoter sequence for efficient repression of divergent noncoding transcription.

**Figure 3.**
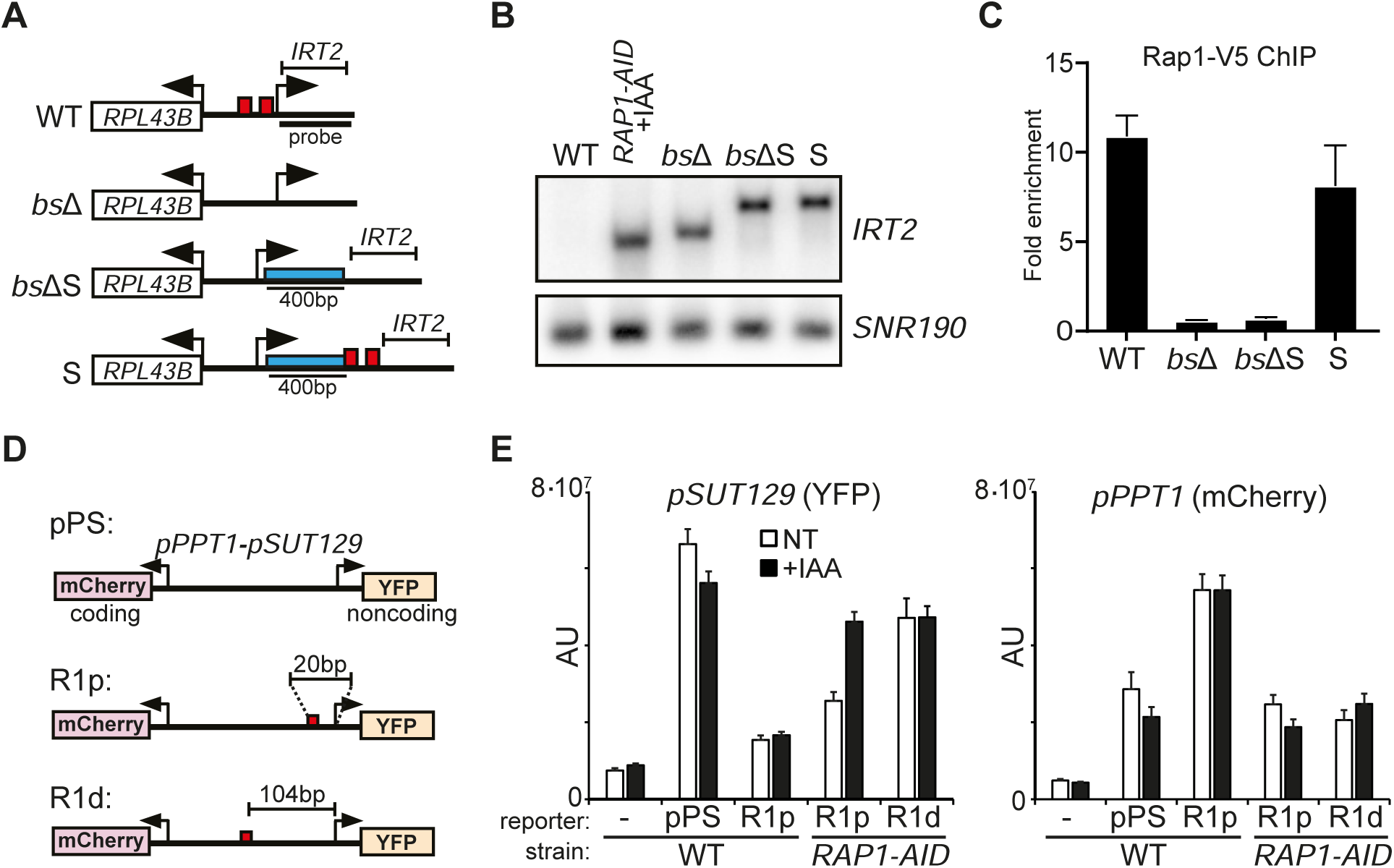
A proximal Rap1 motif is required and sufficient to repress divergent transcription. **(A)** Schematic diagram of mutants used to determine the effect of a distal Rap1 motif on divergent noncoding transcription. Shown are strains with *RPL43B* promoter Rap1 sites deleted (*bsΔ*, FW3440), Rap1 sites deleted and containing a spacer sequence (400bp) (*bs*ΔS, FW3920), or wild-type Rap1 sites and containing a spacer (S, FW3922). Blue bar represents the spacer sequence and red boxes represent Rap1 binding sites. **(B)** *IRT2* expression in strains described A. Wild-type (WT, FW627) and Rap1 depleted cells (*RAP1-AID* +IAA, FW3877) were also included in the analysis. Samples were collected from cells grown to exponential phase. Lanes 1-3 show the same data as Figure 1E. Northern blot membranes were probed for *IRT2* and *SNR190*. **(C)** Rap1 binding in *RPL43B* locus mutants described in A, measured by chromatin immunoprecipitation. Rap1 was tagged with V5 epitope (FW4732, FW4734, FW4737, and FW4735). Signals were normalized over a primer pair directed against the 3’ end of the *ACT1* gene. The mean value + SEM is plotted (n = 3). **(D)** Diagram of fluorescent reporter constructs. The *PPT1-SUT129* divergent promoter construct expressing mCherry in the *PPT1* direction and YFP in the *SUT129* direction (pPS) was described previously (Marquardt et al., 2014). The Rap1 sites from the *RPL43B* promoter were integrated at a proximal (R1p, 20 bp), or distal (R1d, 104 bp) position to the TSS of *SUT129*. The Rap1 sites were cloned in the forward direction relative to mCherry. **(E)** Ectopic repression of divergent noncoding transcription by Rap1. For the analyses we used: wild type cells harboring no reporter (FW627), pPS control reporter (FW6407), or R1p (FW6895), and *RAP1-AID* cells containing R1p (FW6206), or R1d (FW6408). Baseline fluorescence was measured from wild type cells harboring no reporter construct (−). Cells were grown to exponential phase and treated with IAA (final concentration 500 μM) or left untreated (NT) for four hours. Cells fixed with formaldehyde and subsequently imaged. For each sample, mean signals corrected for background (AU, arbitrary units) are plotted with error bars representing 95% confidence intervals (n = 50 cells per sample).

Next, we determined whether the Rap1 binding site on its own is sufficient to repress divergent transcription. We integrated Rap1 motifs into a fluorescent reporter construct that harbours a divergent promoter transcribing *PPT1* in the coding direction and *SUT129* in the noncoding direction (pPS) (Figure 3D and Figure S3A) (Marquardt et al., 2014). Cells harbouring a Rap1 motif proximal to the *SUT129* promoter (R1p) showed decreased YFP levels, while *PPT1* (mCherry) activity increased (Figure 3E, left panel). *SUT129* promoter (R1p) activity increased to match control plasmid (pPS) levels upon Rap1 depletion *(RAP1-AID* +IAA). The repression of *SUT129* by Rap1 was not dependent on transcription regulation in the coding direction because in *RAP1-AID* (IAA or NT) cells the *PPT1* signal closely matched the WT reporter (Figure 3E, right panel). Furthermore, the results were comparable when we used a reporter with Rap1 motifs in the reverse orientation (R1prv) (Figure S3B). Finally, we found that a more distal Rap1 binding site (R1d) to *SUT129* showed comparable YFP levels to the WT reporter, and Rap1 depletion also had little effect (Figure 3E). These data demonstrate that the Rap1 motif, independent of its orientation, is sufficient to repress divergent noncoding transcription when located near the cryptic promoter sequences.

### TSS mapping of Rap1 repressed divergent noncoding transcripts

Our data revealed that close proximity of the Rap1 motif to the cryptic promoter is essential for repressing divergent noncoding transcription. To investigate the relationship between Rap1 motifs and cryptic promoters at a genome-wide scale, we mapped transcription start sites by sequencing (TSS-seq) in wild-type and Rap1 depleted cells *(RAP1-AID* +IAA) (Figure S4A and S4B). At the *RPL43B* promoter, which harbours two Rap1 binding sites, a cluster of multiple TSS was detected in a region of 35 bp up-and downstream of the Rap1 motifs in Rap1 depleted cells and to a lesser extent in wild-type cells (Figure 4A, left panel). The signals are unlikely to originate from abortive RNA polymerase II initiation because the TSS-seq procedure isolates polyadenylated and capped RNAs. At the *RPL40B* promoter, multiple *iMLP1* TSSs in a region of 23 bp were detected directly upstream of the Rap1 binding site in the *RPL40B* promoter, and the TSS-seq signals increased upon Rap1 depletion (Figure 4A, right panel). Conversely, the *MLP1* protein coding TSS signal was decreased in Rap1 depleted cells supporting our earlier observation that Mlp1 protein levels are reduced in cells mis-expressing *iMLP1*.

**Figure 4.**
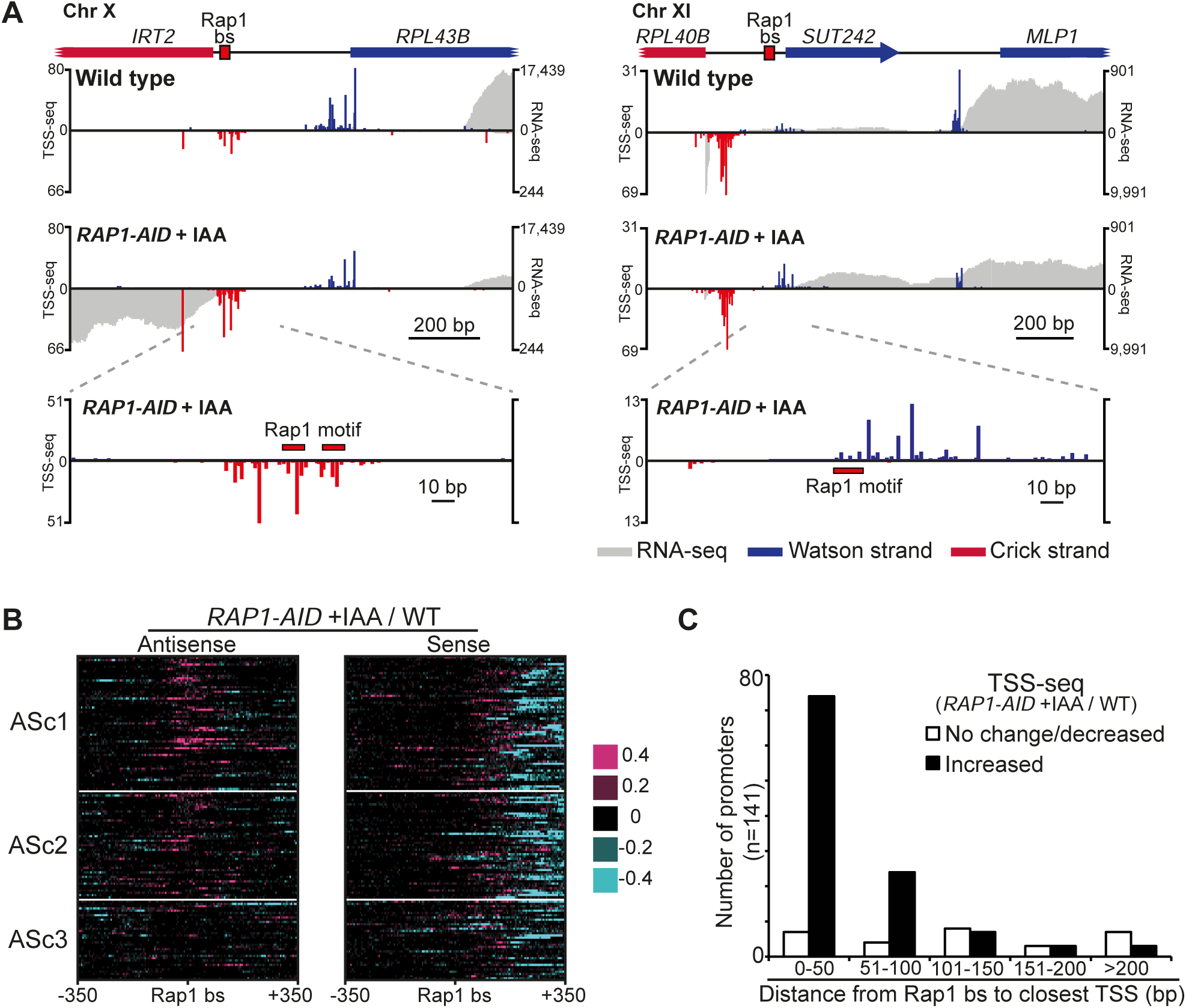
Rap1 represses divergent transcription initiation close to its binding site. **(A)** Genome browser tracks displaying transcription start sites (TSSs) at the *RPL43B* and *RPL40B* loci. Data from wild-type (FW629) and *RAP1-AID* strains (2 hours after IAA treatment, FW3877) are shown. TSS-seq signal per million reads (Watson (W, Blue) and Crick (C, Red)) is plotted for the first mapped nucleotide. Total RNA-Seq tracks (gray) are shown for the Watson strand (W, above) and Crick strand (C, below). **(B)** Difference heat map showing increased divergent noncoding transcription near Rap1 binding sites. Difference ratios for bins of 5 bp around 141 promoter Rap1 binding sites were calculated, comparing the normalized TSS-seq signal after Rap1 depletion (*RAP1-AID +IAA*, FW3877) versus wild type control (WT, FW629). Pink coloured regions represent TSS-seq signal that is higher in the *RAP1-AID* +IAA condition versus WT, and cyan coloured regions represent signal that is lower. Promoters for both antisense and sense plots are clustered and ordered as in Figure 2F, based on RNA-seq signals on the antisense strand. **(C)** Distribution of TSSs near promoter Rap1 binding sites and response to Rap1 depletion from data described in A and B. The distance from the Rap1 binding site (n = 141) to the closest TSS was measured and their distribution is plotted in bins of 50 bp. For the 109 TSSs within 100 bp of Rap1 binding sites, we observed an increased TSS-seq signal for 96 TSSs (88%) and no change/decreased signal for 13 TSSs (12%).

Next, we computed the changes in TSS signal between wild-type and Rap1 depleted cells. The TSS-seq data matched the RNA-seq data well. Clusters 1 and 2 for the antisense orientation (ASc1 and ASc2) displayed increased TSS signals around the Rap1 binding sites, whereas there were fewer differences in cluster 3 (ASc3) (Figure 2F and Figure 4B, antisense). As expected, TSS signals decreased in the sense direction downstream of the Rap1 sites in Rap1 depleted cells because coding gene expression was reduced (Figure 4B, sense). Interestingly, sequences directly upstream of the canonical coding transcript TSSs displayed increased TSS signals in the sense direction, suggesting that Rap1 is also important for TSS selection. Indeed, it has been proposed that Rap1 regulates TSS selection through multiple mechanisms (Kasahara et al., 2011; Reja et al., 2015). Finally, we determined the direction and position of the closest TSS to the Rap1 binding site (Figure 4C). We found that the majority of Rap1 regulated promoters we examined contained an antisense TSS (82% antisense versus 18% sense direction) as the nearest one to the Rap1 binding site. Approximately 50% of the promoters displayed increased TSS signals within 50 bp of the Rap1 motif in Rap1 depleted cells (Figure 4C). Taken together, our analysis demonstrates that Rap1 represses initiation of divergent transcription close to its promoter regulatory elements.

### The Rap1 carboxy-terminal domain contributes to repressing divergent transcription

How does Rap1 repress divergent transcription? One possibility is that a specific function of Rap1 is required. Distinct domains of Rap1 are important for exerting different functions in gene repression and activation (Azad and Tomar, 2016; Shore, 1994). To examine whether repression of divergent transcription requires a specific domain of Rap1, we generated deletions in the N- and C-terminal domains of Rap1 without disrupting the DNA binding domain (Figure 5A). The Rap1 fragments were expressed in *RAP1-AID* cells (Figure S5A). As expected, full-length Rap1 (FL, 1-827) was able to maintain repression of *IRT2* and *iMLP1* expression upon Rap1 depletion (*Rap1-AID* +IAA), whereas the empty vector control (EV) displayed divergent transcription (Figure 5B). A deletion of the N-terminus (ΔN, 339-827) was able to rescue Rap1 depletion. Cells harbouring deletions in the C-terminus (ΔC, 1-599) or N-and C-terminus (ΔN ΔC, 339-599) displayed expression of *IRT2* and *iMLP1*. It is worth noting that the expression of *IRT2* and *iMLP1* in ΔNΔC was decreased to ~70% of EV, indicating that the Rap1 DNA binding domain represses divergent transcription to some extent (Figure S5B). Thus the N-terminus, but not the C-terminus, of Rap1 is dispensable for repression of divergent transcription.

**Figure 5.**
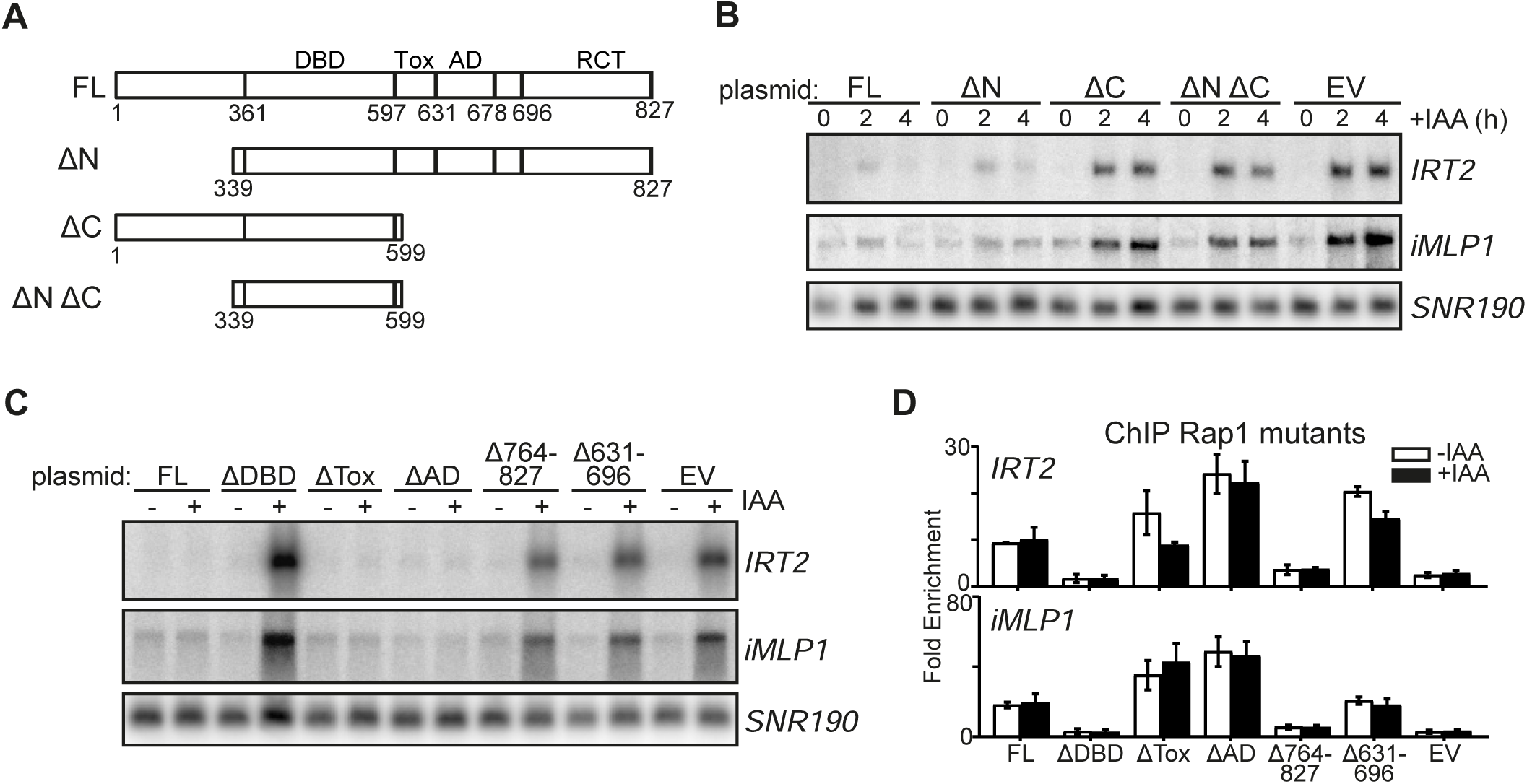
The Rap1 C-terminal domain contributes to repression of divergent noncoding transcription. **(A)** Schematic diagram of domains in Rap1 and mutants with truncations in Rap1. The residues of the DNA-binding domain (DBD), toxicity domain (Tox), activation domain (AD), and Rap1 C-terminal interacting domain (RCT) are displayed. **(B)** *IRT2* and *iMLP1* expression in *RAP1-AID* cells expressing Rap1 truncation mutants. *RAP1-AID* (FW3877) cells also expressing full length Rap1 (FL) (FW4869), ΔN (FW4871), ΔC (FW4874), ΔN ΔC (FW4876), or an empty vector (EV) (FW4878) were grown to exponential phase and treated with IAA (500 μM). **(C)** *IRT2* and *iMLP1* expression in different Rap1 domain mutants. For the analysis we used *RAP1-AID* cells expressing Rap1 full length (FL) (FW4948), DNA binding domain deletion (ΔDBD) (FW4950), toxicity domain deletion (ΔTox) (FW4952), activation domain deletion (ΔAD) (FW4954), deletion of residues 764 to 827 (Δ764-827) (FW4958), deletion of residues 631 to 696 (Δ631-696) (FW4960), or empty vector (EV) (FW4878). **(D)** ChIP of Rap1 mutants described in E, at *RPL43B (IRT2)* and *RPL40B (iMLP1)* promoters. Similar mutants as in E, except that a V5 tag was introduced at the C-terminus and the depletion was performed using a *RAP1-AID-MYC* allele (FW5420, FW5393, FW5394, FW5424, FW5395, FW5396 and FW5399). The mean fold enrichment value ± SEM is plotted (n = 3).

Important functions of the C-terminus of Rap1 are exerted by the silencing domain, the activation domain (AD), and the toxicity domain (Tox) (Freeman et al., 1995; Garbett et al., 2007; Johnson and Weil, 2017; Kurtz and Shore, 1991; Layer et al., 2010; Sussel and Shore, 1991). We assessed whether Rap1 constructs with different domain deletions could repress divergent transcription (Figure 5A) (Layer et al., 2010). The Rap1 domain mutants were expressed in Rap1 depleted cells (Figure S5C-E). We found that Rap1ΔTox and Rap1ΔAD did not affect *IRT2* and *iMLP1* repression, whereas mutants lacking the DNA binding domain (Rap1ΔDBD), the silencing domain (Rap1Δ764-827), or the activation domain plus an adjacent sequence (Rap1Δ631-696) failed to repress *IRT2* and *iMLP1* (Figure 5C). Except for Rap1ΔDBD and Rap1Δ764-827, the Rap1 C-terminal mutants associated at the *RPL43B (IRT2)* and *RPL40B (iMLP1)* promoters (Figure 5D and Figure S5D). Given that Rap1Δ764-827 was not able to bind to Rap1 sequence elements, we examined whether different point and patch mutations in the Rap1 silencing domain, already characterized for telomere regulation and hidden mating-type loci silencing, affected repression of *IRT2* expression (Feeser and Wolberger, 2008). A summary of the data is listed in Table S1. We found that none of the mutants caused a significant increase in *IRT2* expression indicating the Rap1 silencing domain is not important for repressing divergent transcription (Table S1). We conclude that part of Rap1 C-terminus, which includes the activation domain but not the silencing domain, contribute to repression of divergent transcription.

### RSC chromatin modeller elicits divergent transcription in the absence of Rap1

Given that the Rap1Δ631-696 mutant displayed divergent transcription but maintained its ability to bind the Rap1 motif, we hypothesized that association of regulators of divergent transcription such as co-repressors or activators may be altered in this mutant. To identify candidate regulators of divergent transcription, we isolated chromatin-bound Rap1 and associated proteins from cells. We affinity-purified V5-tagged Rap1 from micrococcal nuclease (MNase) solubilized chromatin and used proteomics mass spectrometry to identify associated proteins (Figure 6A) (van Werven et al., 2008). For the analyses we used full-length Rap1 (Rap1-FL), Rap1ΔAD, Rap1Δ631-696 and an empty vector control (Figure S6A). First, we determined the wild-type Rap1 chromatin mediated interactome by comparing Rap1-FL to the empty vector control (Figure 6B). Several proteins from complexes known to interact with Rap1 were enriched in Rap1-FL compared to the control, e.g. TAFs, telomere related proteins, and nuclear pore complex proteins (NPCs) (Layer et al., 2010; Shi et al., 2013; Van de Vosse et al., 2013). In addition, multiple subunits of the chromatin remodeller RSC (12 out of 17) were enriched in Rap1-FL compared to the empty vector (Figure 6B). Gene Ontology (GO) analysis revealed that 72 out of 276 enriched proteins were involved in RNA polymerase II transcription, and 66 proteins were involved in chromatin organization (Figure 6C and Figure S6B). Next, we specifically searched for interacting proteins that showed differential enrichment between Rap1Δ631-696 and Rap1-FL, but which were not altered in Rap1ΔAD. We found that all identified subunits (12 out of 17) of RSC were enriched in Rap1Δ631-696, but not in Rap1ΔAD, compared to Rap1-FL (Figure 6D). These data suggest that RSC could play a role in controlling divergent transcription at Rap1 regulated gene promoters.

**Figure 6.**
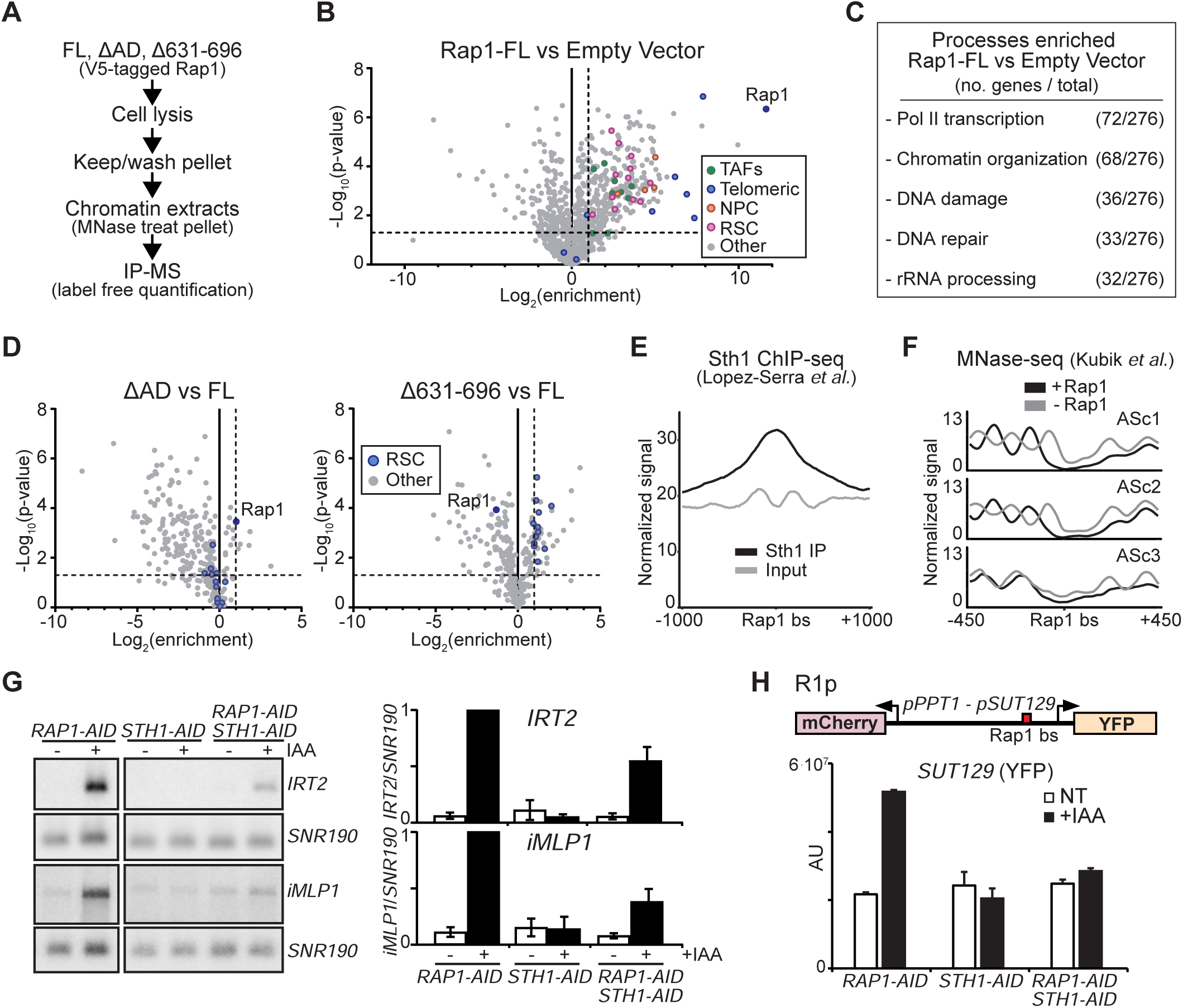
RSC chromatin modeller elicits divergent transcription in the absence of Rap1. **(A)** Experimental scheme used to identify proteins interacting with chromatin bound Rap1. Chromatin was solubilized by micrococcal nuclease (MNase) digestion. V5-tagged Rap1 constructs for FL (FW5420), ΔAD (FW5424), Δ631-696 (FW5396) and an empty vector control (FW5399), were affinity purified and processed for LC-MS label free quantification (LFQ). **(B)** Volcano plot displaying differences in protein enrichment for Rap1-V5 (FL) versus empty vector control. The enrichment (Log_2_) versus *p*-value (unpaired two-sample *t*-test, −Log_10_ scale) for n = 916 identified proteins is displayed. Highlighted proteins: bait protein (Rap1, dark blue), TFIID TAFs (green), telomeric proteins (blue), NPC components (orange), and RSC components (purple). Horizontal dashed line corresponds to 1.303 (*p* = 0.05), and vertical dashed line corresponds to 2-fold enrichment. (C) Yeast GO-Slim Process Analysis of proteins enriched from data described in B. **(D)** Volcano plots of ΔAD versus full length Rap1 (left) and Δ631-696 versus full length Rap1 (right). Proteins that were in enriched in full-length versus empty vector control (n = 289 proteins) as described in 6B are plotted. Highlighted proteins: bait protein (Rap1, dark blue), RSC complex components (blue), NPC components (orange). **(E)** Metagene plots of Sth1 ChIP-seq data for Rap1 regulated promoters (n = 141), centered on Rap1 binding sites (bs) (Lopez-Serra et al., 2014). (F) Metagene plots of MNase-seq data (Kubik et al., 2015) for three clusters of promoters based on divergent noncoding transcription (ASc1, ASc2, ASc3) as described in Figure 2F. Displayed are the signals in the presence (black) or absence (gray) of Rap1. **(G)** *IRT2* and *iMLP1* expression in cells co-depleted for Sth1 and Rap1. Cells harbouring *RAP1-AID* (FW3877), *STH1-AID* (FW6032), and *RAP1-AID STH1-AID* (FW6231) alleles were grown to exponential phase, and samples were collected before (−) and 2 hours after (+) treatment with IAA (500 μM). Membranes were probed for *IRT2, iMLP1* and *SNR190* (left). Quantification of *IRT2* and *iMLP1* expression (right). The signal was normalized over *SNR190*. The normalized signal *IRT2* or *iMLP1* expression in Rap1 depleted cells (*RAP1-AID* +IAA) was set to 1. Mean values ±SEM are plotted (n = 3). **(H)** *SUT129* promoter activity is suppressed by co-depletion of RSC and Rap1. *RAP1-AID* (FW6206), *STH1-AID* (FW6218), and *RAP1-AID STH1-AID* (FW6433) cells harbouring the R1p construct were treated with IAA (500 μM) or left untreated (NT). *SUT129* activity (YFP) of the R1p reporter construct was measured as described in Figure 3G. For each sample, mean signals corrected for background (AU, arbitrary units) are plotted plus 95% confidence intervals for n = 50 cells.

RSC is an ATP-dependent chromatin remodelling complex and an important regulator of nucleosome organization (Cairns et al., 1996). In particular, RSC plays a critical function at promoters by generating a nucleosome-depleted region (NDR), thereby facilitating activation of gene expression (Badis et al., 2008; Hartley and Madhani, 2009; Lorch et al., 2011; Parnell et al., 2008). The ATPase subunit of RSC, Sth1, binds near promoter Rap1 binding sites, supporting our observation that RSC can interact with chromatin bound Rap1 (Figure 6E and Figure S6C) (Lopez-Serra et al., 2014; Parnell et al., 2015). The RSC complex interacts with nucleosomes and DNA directly, and polyA and GC-rich motifs direct its recruitment and action (Floer et al., 2010; Krietenstein et al., 2016; Kubik et al., 2015). Thus RSC may not require Rap1 for promoter association, which is in line with observation that a narrow NDR is maintained at Rap1-regulated gene promoters in the absence of Rap1 (Figure 6F and Figure S6D) (Kubik et al., 2015). It is worth noting that for the clusters with high levels of divergent transcription (ASc1 and ASc2) nucleosomes are highly organized directly upstream of the Rap1 motif, which is likely caused by transcription coupled chromatin remodelling (Figure 6F and Figure S6D) (Venkatesh and Workman, 2015). Thus, RSC binds to Rap1 regulated gene promoters, and may be important for maintaining open chromatin independent of Rap1. A recent study showed that RSC binding to promoters is independent of Rap1 and other pioneer transcription factors (Kubik et al., 2018).

Given the role of RSC in regulating nucleosome positioning, and our observation that association of RSC with the Rap1Δ631-696 mutant was increased, we hypothesized that RSC promotes divergent transcription in the absence of Rap1. To test this, we depleted Sth1 together with Rap1 (Figure S6E). Depleting Sth1 *(STH-AID* +IAA) by itself had no effect on *IRT2* and *iMLP1* expression (Figure 6G). When Rap1 and Sth1 were co-depleted, *IRT2* and *iMLP1* expression was greatly reduced compared to Rap1 depletion alone (Figure 6G and Figure S6F). Depleting RSC also supressed divergent transcription when we used the *PPT1/SUT129* reporter plasmid harbouring proximal Rap1 sites (R1p, Figure 6H and Figure S6G). Whereas Sth1 depletion had no effect on *SUT129* promoter activity, co-depletion of Sth1 with Rap1 suppressed the increased *SUT129* signal observed in Rap1 depleted cells. Thus, RSC promotes divergent noncoding transcription in the absence of Rap1. We propose that Rap1 is positioned to repress divergent noncoding transcription elicited by RSC and thereby restricts RSC to stimulate productive transcription in the protein-coding direction.

### Chromatin regulators control divergent transcription in a manner distinct from Rap1

Chromatin remodellers and histone modifying enzymes play essential roles in repressing noncoding transcription (Venkatesh and Workman, 2015). We hypothesized that Rap1 mediated repression of divergent transcription could be facilitated or mediated by specific chromatin regulators. To identify additional repressors of divergent transcription at Rap1 regulated gene promoters, we measured *IRT2* and *iMLP1* expression levels in different gene deletion and depletion strains. Specifically, we selected genes that are (1) involved in cryptic or divergent transcription (e.g. Set2, Set3, and Spt16), (2) known to interact with Rap1 (e.g. Sir2, Rif1, and Rif2), and (3) other chromatin and transcription regulators. We assessed the expression of *IRT2* and *iMLP1* in 62 deletion and 4 depletion strains (Table S2). Fourteen mutants displayed increased *iMLP1* expression. Only depletion of Spt16 (*SPT16-AID* +IAA) increased *IRT2* expression. When we compared the *iMLP1* expression patterns in our data to a published dataset we found that five mutants overlapped, which we decided to study further (van Bakel et al., 2013). These were: (1) putative histone deacetylase Spt10, (2) transcription factor Spt21, (3) CAF-1 chromatin assembly complex component Rlf2, (4) chromatin remodeller and elongation factor Spt6, and (5) FACT complex component Spt16. All candidates have known roles in repression of divergent or cryptic transcription (Cheung et al., 2008; DeGennaro et al., 2013; Jeronimo et al., 2015; Marquardt et al., 2014). We performed RNA-seq using gene deletion and depletion alleles for all five chromatin regulators and observed increased expression within Rap1 regulated promoters (Figure 7A and Figure S7A-B). These data show that multiple chromatin regulators contribute to repression of divergent noncoding transcription at Rap1 regulated gene promoters.

**Figure 7.**
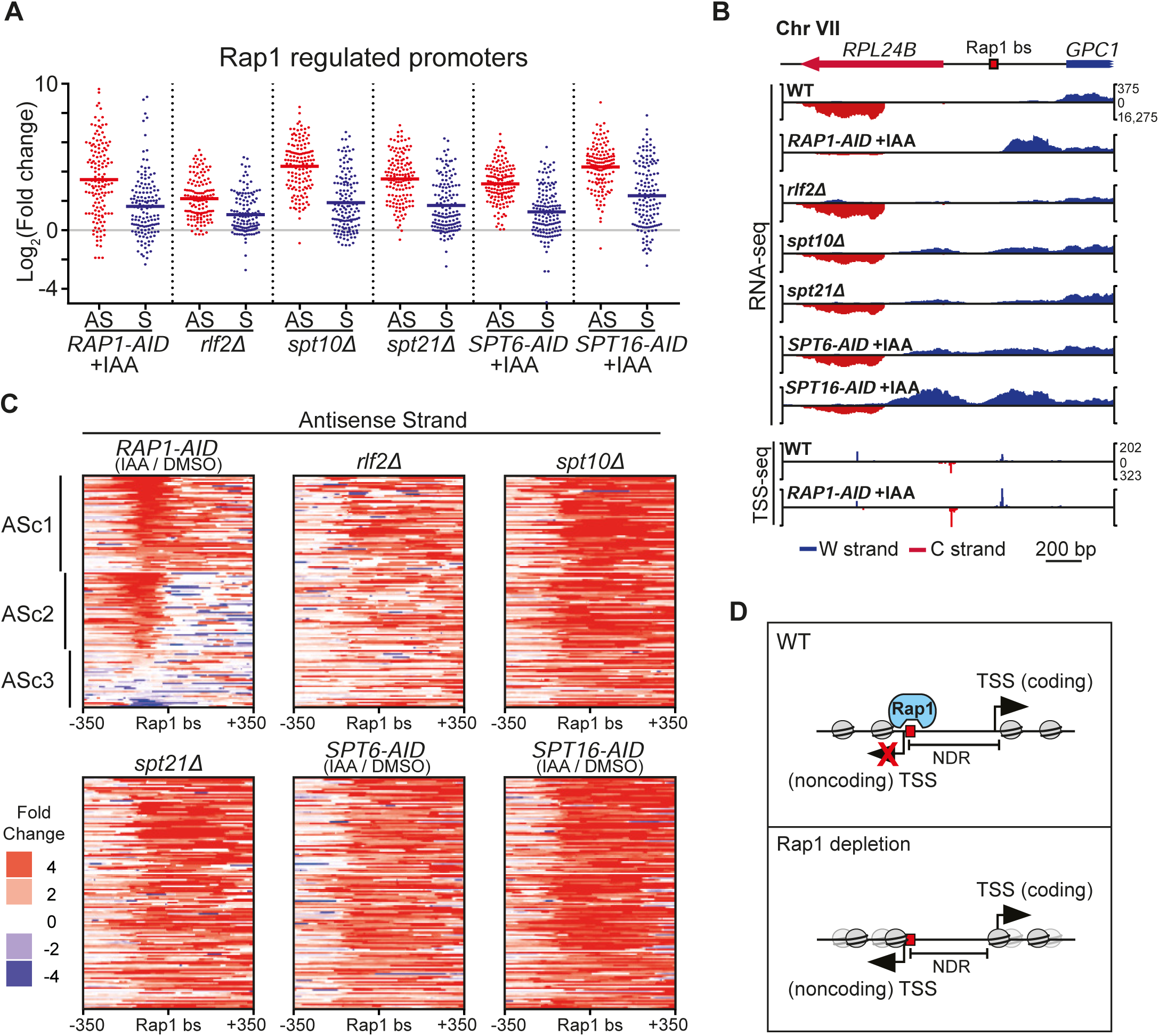
Chromatin regulators control divergent transcription in a manner distinct from Rap1. **(A)** Scatter plots showing the changes in RNA expression around promoter Rap1 sites. Wild type (FW629), *spt10Δ* (FW5543), *spt21Δ* (FW5547), and *rlf2Δ* (FW5609) cells were grown to exponential phase and samples were taken for total RNA-seq. In addition, samples were collected from *RAP1-AID* (FW3877), *SPT6-AID* (FW5555), and *SPT16-AID* (FW5559) cells that were treated for two hours with IAA (500 μM) or DMSO. Changes in RNA expression were calculated for n =141 promoter Rap1 sites ±100 bp for antisense and sense strands. Horizontal red or blue bars indicate mean values. **(B)** Data from A displayed in genome browser tracks for the *RPL24B* locus. The corresponding TSS-seq data described in Figure 4 are also displayed. **(C)** Heat maps showing the RNA expression differences around Rap1 binding sites for data described in A and B. Data for all plots are clustered and ordered according to antisense strand signals from *RAP1-AID* cells, as described in Figure 2F. **(D)** A model for transcription factor mediated repression of divergent noncoding transcription. When Rap1 is bound at its motif (red boxes), coding direction transcription is activated and initiates from the coding direction TSS (black arrows) but initiation from the divergent noncoding TSS is repressed. When Rap1 is absent, transcription in the divergent noncoding direction occurs.

Next, we examined whether chromatin regulators (Rlf2, Spt10, Spt21, Spt6, and Spt16) mediate the repression of divergent transcription by Rap1. We found little overlap between Rap1 repressed divergent transcripts and transcripts repressed by the five chromatin regulators. To illustrate, the *RPL24B* and *RPL40B* promoters showed antisense transcription downstream of the Rap1 motif nearer to or within the coding gene in *rlf2Δ, spt10Δ, spt21Δ* cells, and in cells depleted for Spt6 and Spt16 (*SPT6-AID* +IAA and *SPT16-AID* +IAA) (Figure 7B and Figure S7C). We also identified promoters *(RPL25* and *RPL43B)* that displayed no detectable divergent transcription in these depletion and deletion mutants, while there was a clear signal in Rap1 depleted cells *(RAP-AID* +IAA) (Figure S7C). We grouped and ordered the data according to the gene clusters identified in RNA-seq from Rap1 depleted cells (Figure 7C and Figure S7D). All five depletion or deletion mutants displayed increased divergent transcription (ASc1-3) which initiated from within gene bodies and downstream of the Rap1 binding sites, but not initiating near Rap1 binding sites as we observed in Rap1 depleted cells (Figure 7C). Taken together, our data suggest that Rap1 acts in concert with chromatin regulators to repress cryptic or divergent transcription, but in a clearly distinct manner that is spatially limited.

## Discussion

Eukaryotic cells use various mechanisms to tightly control gene expression and limit accumulation of unwanted transcripts. Transcriptionally active promoters are a major source of pervasive transcription. Here, we described how highly expressed coding gene promoters limit divergent noncoding transcription in yeast. We identified a surprising role for the pioneer transcription factor Rap1. We found that Rap1 represses divergent noncoding transcription at its binding motif and adjacent sequences. Our data demonstrate the first example of a sequence specific transcription factor that can prevent regulatory sequences from producing aberrant transcripts. Our study defines a novel mechanism for providing directionality towards productive transcription.

### Mechanism of Rap1 mediated repression of divergent transcription

Limiting pervasive transcription is a key feature of gene regulation. We identified a transcription factor mediated mechanism for limiting pervasive transcription. Several lines of evidence indicate that Rap1 specifically represses divergent noncoding transcription, which is uncoupled from transcription regulation in the coding direction. First, Rap1 represses divergent transcription near its binding site, typically within 50 base pairs of the Rap1 motif. Second, abrogating other transcription factors important for ribosomal protein gene expression did not affect divergent transcription, supporting a specific function for Rap1. Third, close proximity of the Rap1 binding site to the cryptic core promoter is essential for repressing divergent transcription. Fourth, the Rap1 binding site ectopically represses divergent noncoding transcription without affecting transcription in the protein-coding direction. Conversely, the activation domain of Rap1, which directs transcription in the protein-coding direction, is not required for repressing divergent transcription (Johnson and Weil, 2017; Layer et al., 2010). Finally, we found that a chromatin assembly factor (Rlf2), regulators of histone gene expression (Spt10 and Spt21), and co-transcriptional chromatin remodellers (Spt6 and Spt16) also repress divergent transcription by mechanisms distinct from Rap1.

Our data show that Rap1 mediated repression of divergent transcription confers promoter directionality. Transcription directionality is shaped by evolution towards protein-coding genes specifically through enrichment of DNA binding protein motifs (Jin et al., 2017). In this context, Rap1 promotes directionality in multiple ways. First, Rap1 recruits cofactors and basal transcription machinery which promote transcription in the coding direction (Garbett et al., 2007; Hall et al., 2006; Marion et al., 2004; Papai et al., 2010; Reja et al., 2015; Rudra et al., 2005; Wade et al., 2004; Zhao et al., 2006). Second, Rap1 asymmetrically occupies the promoter NDR at the 5’ end, where it represses the divergent core promoter (Figure 7C) (Kubik et al., 2015; Reja et al., 2015). Core promoters are intrinsically directional (Duttke et al., 2015), and two independent PICs initiate divergent transcription at mRNA-ncRNA pairs in yeast (Rhee and Pugh, 2012). Hence, repressing transcription initiation at the core promoter in the antisense direction promotes overall promoter directionality. In mammalian cells, some pioneer transcription factors also open chromatin asymmetrically (Sherwood et al., 2014) – suggesting that repression of divergent transcription by sequence-specific transcription factors could be conserved across species.

Our findings have functional implications on the positioning and organization of TSSs and transcription factor binding sites at promoters. In yeast, an optimal distance between the upstream activating sequence (UAS) and core promoter is important for productive transcription (Dobi and Winston, 2007). Our data suggest that upstream regulatory elements or transcription factor binding sites should not overlap with core promoters to avoid concurrent steric interference. A regulatory element too far or too close to the core promoter will affect coding gene expression. The position of regulatory elements or transcription factor binding sites within NDR also has functional consequences for coding and divergent transcription. Some transcription factors associate in the middle of the NDR, which may promote divergent or bidirectional transcription (Sherwood et al., 2014). In the case of Rap1 and other transcription factors, the regulatory elements are positioned asymmetrically at the 5’ border of the NDR (Sherwood et al., 2014). This may be a requirement of transcription factors that limit divergent transcription. It has been proposed that DNA sequences and protein co-evolved to promote productive directional transcription (Jin et al., 2017). Our study of Rap1 illustrates a mechanism by which enrichment of DNA sequences and transcription factors towards asymmetric promoter binding effectively limits divergent transcription. Overall, further investigation is required to fully understand the mechanistic details by which regulatory elements, TSSs, and NDRs control gene expression and divergent transcription.

How does Rap1 repress divergent noncoding transcription? We found no evidence that the Rap1 silencing function is important. It also seems unlikely that the Rap1 roadblock function is important for repressing divergent transcription (Candelli et al., 2018; Yarrington et al., 2012). Typically, the Rap1 roadblock acts as a failsafe mechanism by terminating transcriptional read-through of upstream coding and noncoding RNAs. While Rap1-mediated repression of divergent transcription is antisense to the coding direction, the Rap1 roadblock terminates elongating polymerases originating upstream in the sense direction towards the coding gene. Many Rap1-repressed divergent transcripts initiate directly upstream of the Rap1 motif. In other words, there is no Rap1 motif downstream from the transcription initiation site of divergent transcripts.

Rap1-mediated repression of divergent transcription shows parallels to prokaryotic operon regulation and certain synthetic transcriptional repression systems. In bacteria, transcriptional repressors bind operon sequences near TSSs and directly prevent recruitment of RNA polymerase through steric hindrance (Browning and Busby, 2004; Lloyd et al., 2001; Rojo, 1999). Similarly, Rap1 mediated repression of divergent transcription could also act through steric hindrance. Like bacterial repressors, Rap1 binds near the TSS of (divergent) core promoters. In addition, our data suggest that Rap1 represses divergent transcription directly because we find no evidence for other contributions from Rap1 cofactors. Our analysis revealed that residues within the C-terminus of Rap1 contribute to repression of divergent transcription. Perhaps, the C-terminal region we identified contributes to steric hindrance, DNA binding affinity or protein stability. It has been suggested that the Rap1 C-terminus can modulate the affinity and binding mode of the DNA binding domain (Feldmann et al., 2015; Feldmann and Galletto, 2014). In eukaryotes, direct steric repression of transcription can also be mediated by CRISPR interference (CRISPRi) and transcription activator-like effector repressors (TALERs) (Gilbert et al., 2013; Li et al., 2015; Qi et al., 2013). Like Rap1, the catalytically inactive dCas9 and TALERs repress gene expression when targeted near TSSs. Taken together, these data suggest a conserved ability of sequence-specific transcription factors to repress transcription initiation by steric hindrance. We propose that steric hindrance by Rap1 prevents Rap1 binding sites and adjacent sequences from initiating divergent transcription (Figure 7D).

### An interplay between Rap1 and RSC controls divergent transcription

Our data show that an interplay between Rap1, the RSC chromatin remodeller, and nucleosomes controls divergent transcription. In Rap1 depleted cells RSC elicits divergent transcription, suggesting that Rap1 restricts RSC activity to stimulate productive transcription in the protein-coding direction only. Recently, it was demonstrated that RSC helps maintain an NDR in the absence of Rap1 (Kubik et al., 2018). Our data suggest that a RSC-dependent NDR contributes to divergent transcription in Rap1 depleted cells. We propose that in wild-type cells, Rap1 occupancy competes locally with the binding of activators of divergent transcription, such as RSC and basal transcription machinery (Figure 7D). Further analysis of specific promoter architectures could elucidate how Rap1, RSC, and other chromatin remodellers control divergent noncoding transcription.

### A model for control of divergent noncoding transcription

We have shown that the pioneer transcription factor Rap1 represses divergent noncoding transcription. Mis-regulation of divergent noncoding transcripts could have negative effects on local or global gene expression, especially in gene-dense genomes such as budding yeast. Decades of work have shown that eukaryotes from yeast to metazoans have adopted important and redundant strategies to limit expression of aberrant noncoding RNAs, including chromatin regulation, transcription initiation and termination, and RNA degradation (Jensen et al., 2013; Porrua and Libri, 2015; Venkatesh and Workman, 2015; Xue et al., 2017). Our findings add a new layer of regulation to the various mechanisms that limit expression of noncoding transcripts. We propose that repression of cryptic transcription nearby regulatory elements could be an evolutionary conserved property of sequence-specific transcription factors and other DNA binding proteins.

## Supplemental Information

Supplemental information contains seven figures (Figure S1-S7), six tables (Table S1-S5), and Materials and Methods.

## Author Contributions

AW and FW conceptualized the study; AW, HP, MC, FM, DF, AS, and FW designed the experiments; AW, MC, FM, DF, and FW performed the experiments; AW, HP, FM, DF, and FW analyzed the data; AW and FW wrote the manuscript with input from all authors. FW supervised the project.

## Acknowledgements

We are grateful to Frank Uhlmann, Jesper Svejstrup and members of the Van Werven lab for their advice and their critical reading of the manuscript. We thank the Crick Advanced Sequencing and Genomics Equipment Park Facilities for experimental support and analysis, Peter Thorpe, Amanda Johnson, Tony Weil, Cynthia Wolberger, Sebastian Marquardt, and Jesper Svejstrup for sharing reagents, and Domenico Libri for sharing unpublished data. This work was supported by the Francis Crick Institute (FC001203), which receives its core funding from Cancer Research UK (FC001203), the UK Medical Research Council (FC001203), and the Wellcome Trust (FC001203). MC is supported by a graduate student fellowship from the Agency for Science, Technology and Research (A*STAR) of Singapore.

